# SopB- and SifA-dependent shaping of the *Salmonella*-containing vacuole proteome in the social amoeba *Dictyostelium discoideum*

**DOI:** 10.1101/2019.12.23.887745

**Authors:** Camila Valenzuela, Magdalena Gil, Ítalo M. Urrutia, Andrea Sabag, Jost Enninga, Carlos A. Santiviago

## Abstract

The ability of *Salmonella* to survive and replicate within mammalian host cells involves the generation of a membranous compartment known as the *Salmonella-*containing vacuole (SCV). *Salmonella* employs a number of effector proteins that are injected into host cells for SCV formation using its type-three secretion systems encoded in SPI-1 and SPI-2 (T3SS_SPI-1_ and T3SS_SPI-2_, respectively). Recently, we reported that *S*. Typhimurium requires T3SS_SPI-1_ and T3SS_SPI-2_ to survive in the model amoeba *Dictyostelium discoideum*. Despite these findings, the involved effector proteins have not been identified yet. Therefore, we evaluated the role of two major *S*. Typhimurium effectors SopB and SifA during *D. discoideum* intracellular niche formation. First, we established that *S*. Typhimurium resides in a vacuolar compartment within *D. discoideum*. Next, we isolated SCVs from amoebae infected with wild type or the Δ*sopB* and Δ*sifA* mutant strains of *S*. Typhimurium, and we characterized the composition of this compartment by quantitative proteomics. This comparative analysis suggests that *S*. Typhimurium requires SopB and SifA to modify the SCV proteome in order to generate a suitable intracellular niche in *D. discoideum*. Accordingly, we observed that SopB and SifA are needed for intracellular survival of *S*. Typhimurium in this organism. Thus, our results provide insight into the mechanisms employed by *Salmonella* to survive intracellularly in phagocytic amoebae.

**Importance:** The molecular mechanisms involved in *Salmonella* survival to predation by phagocytic amoebae, such as *D. discoideum*, remains poorly understood. Although we established that *S*. Typhimurium requires two specialized type-three secretion systems to survive in *D. discoideum*, no effector protein has been implicated in this process so far. Here, we confirmed the presence of a membrane-bound compartment containing *S*. Typhimurium in *D. discoideum*, and purified the *D. discoideum* SCV to characterize the associated proteome. In doing so, we established a key role for effector proteins SopB and SifA in remodeling the protein content of the SCV that ultimately allow the intracellular survival of *S*. Typhimurium in *D. discoideum*. We also discuss similarities and differences with the proteomes of the human SCV. These findings contribute to unravel the mechanisms used by *Salmonella* to survive in the environment exploiting phagocytic amoebae as a reservoir.

## Introduction

Bacteria from the *Salmonella* genus infect warm-blooded animals targeting the gastrointestinal tract. Particular serotypes, such as *Salmonella enterica* serovar Typhimurium (*S*. Typhimurium), represent major leading causes of gastroenteritis in humans in developing countries (1). Overall, *Salmonella* infections account for over 150,000 deaths annually in the world, most of them associated with foodborne infections (1).

Among the essential genes required for *Salmonella* virulence, pathogenicity islands SPI-1 and SPI-2 encode two independent type three secretion systems (T3SS). These systems are macromolecular structures used to inject effector proteins directly into targeted host cells, and they play major roles during successive steps of the infection cycle. The T3SS encoded in SPI-1 (T3SS_SPI-1_) and its cognate effectors are required during the intestinal phase of the infection, allowing the invasion of epithelial cells through the reorganization of the actin cytoskeleton at the bacterial-host contact sites inducing bacterial endocytosis by these non-phagocytic cells (2, 3). In addition, after crossing the epithelial barrier *Salmonella* may use the T3SS_SPI-1_ to enter into phagocytic cells residing in the sub-epithelium, such as macrophages, neutrophils and dendritic cells. The T3SS encoded in SPI-2 (T3SS_SPI-2_) is expressed upon internalization within different host cell types. This system and its cognate effectors enable *Salmonella* to avoid phagosome-lysosome fusion and to generate the *Salmonella*-containing vacuole (SCV). Within this membrane-bound compartment the pathogen can survive and replicate (4–8).

One understudied aspect of *Salmonella* biology is its survival in the environment, where the pathogen spends an important part of its life cycle. In this niche, *Salmonella* is exposed to predation by protozoa such as amoebas. These eukaryotic microorganisms are phagocytic cells that feed on bacteria. Multiple studies have demonstrated that different *Salmonella* serotypes are able to survive within a number of protozoa genera, such as *Acanthamoeba*, *Tetramitus*, *Naegleria*, *Hartmannella* and *Tetrahymena* (9–16). In the case of other intracellular pathogens, such as *Legionella pneumophila*, the mechanisms to survive and replicate inside amoebae have been studied in some detail (17, 18). Moreover, as in the case of *Salmonella*, *L. pneumophila* injects effector proteins to subvert physiological processes in the host cell in order to survive within amoebae and human macrophages (19, 20). This suggests that the molecular weapons used by intracellular bacterial pathogens have evolved through early interactions with environmental phagocytic organisms. Thus, understanding the mechanisms used by *Salmonella* to survive intracellularly in amoebae will provide insights into how these bacteria acquired the ability to colonize and survive within phagocytic cells, including the macrophages of a given animal host.

Recently we started characterizing the interaction of *Salmonella* with the social amoeba *Dictyostelium discoideum* as this organism has proven to be a powerful model to study host-pathogen interaction for several bacterial pathogens (18, 21–23). Early reports from different groups presented some discrepancies regarding the use of *D. discoideum* as an appropriate host model for *Salmonella* (24, 25). However, we and other authors have recently described that *S*. Typhimurium is able to survive intracellularly (and proliferate at later times of infection) in *D. discoideum*, and that inactivation of genes encoding relevant virulence factors in other models (including T3SS_SPI-1_ and T3SS_SPI-2_) generates defects in intracellular survival in this organism (26–29). These observations support the use of *D. discoideum* as a suitable model to study the processes involved in the intracellular survival of *S*. Typhimurium.

Effectors such as SopB (secreted by T3SS_SPI-1_) and SifA (secreted by T3SS_SPI-2_) have been characterized for their abilities in modifying the SCV composition. In fact, both proteins are major effectors involved in the biogenesis and maturation of the SCV in eukaryotic cells (30). SopB is a phosphatidylinositol phosphatase involved in reducing the levels of PI(4, 5)P_2_ in the nascent SCV and in modifying the repertoire of proteins that interact with this compartment, in particular Rab5 and its cognate PI3K, Vps34 (31, 32). Furthermore, SopB is involved in the recruitment of Rab7 resulting in the accumulation of LAMP1, vATPase and other late endosome markers at the SCV (33–36). On the other hand, SifA is involved in the maintenance of the SCV membrane integrity by its interaction with other *Salmonella* effectors (37). SifA also interacts with the molecular motor kinesin using SKIP as adaptor protein. This interaction is crucial to allow the formation of a network of tubular structures known as the *Salmonella*-induced filaments (SIFs) (38, 39).

In this study, we established that *S*. Typhimurium resides in a vacuolar compartment within *D. discoideum*. SCVs isolated from amoebae infected with wild type (WT) and mutant strains of *S*. Typhimurium were subjected to quantitative proteomics to characterize the composition of this compartment. Comparative analysis of the data suggests that effectors SopB and SifA are involved in the modification of the SCV proteome to ensure a suitable niche for *S*. Typhimurium within *D. discoideum*. Consistently, we established that *S*. Typhimurium requires both SopB and SifA to survive intracellularly in this organism. Overall, our results contribute to understand the mechanisms used by *Salmonella* to survive predation by phagocytic amoebae that may act as reservoirs for this pathogen in the environment.

## Results

### *S*. Typhimurium resides in a vacuolar compartment within infected *D. discoideum*

We evaluated the presence of a vacuolar compartment containing *S*. Typhimurium in *D. discoideum*. For this purpose, infection assays were performed using a reporter *D. discoideum* strain expressing the VatM subunit of the vacuolar ATPase fused to GFP (VatM-GFP). This allowed the visualization of VatM^+^ vacuolar compartments by confocal microscopy. Of note, VatM has been shown to be present at the membrane of different vacuolar compartments in *D. discoideum*, including phagosomes, endosomes, lysosomes and the contractile vacuole (40), and it is one of the proteins recruited to the SCV in other cellular models (41). For our experiments, we used a WT strain of *S*. Typhimurium constitutively expressing fluorescent mCherry (42). We observed that most of the infected amoebae presented VatM+ structures surrounding intracellular bacteria at different times post infection. Representative images of infected amoebae with the typical VatM-GFP staining in the surrounding of red fluorescent bacteria obtained at 3 and 4,5 h post infection are shown in **Supplementary Figure S1A** and **Figure 1**, respectively. We quantified the co-localization between VatM^+^ vacuoles and *S*. Typhimurium and our results indicate that 93% of the intracellular bacteria in infected amoebae are surrounded by the VatM-GFP signal (**Supplementary Figure S1B**). These observations confirm the existence of a VatM^+^ vacuolar compartment containing *S*. Typhimurium within infected *D. discoideum*.

**Figure 1.**
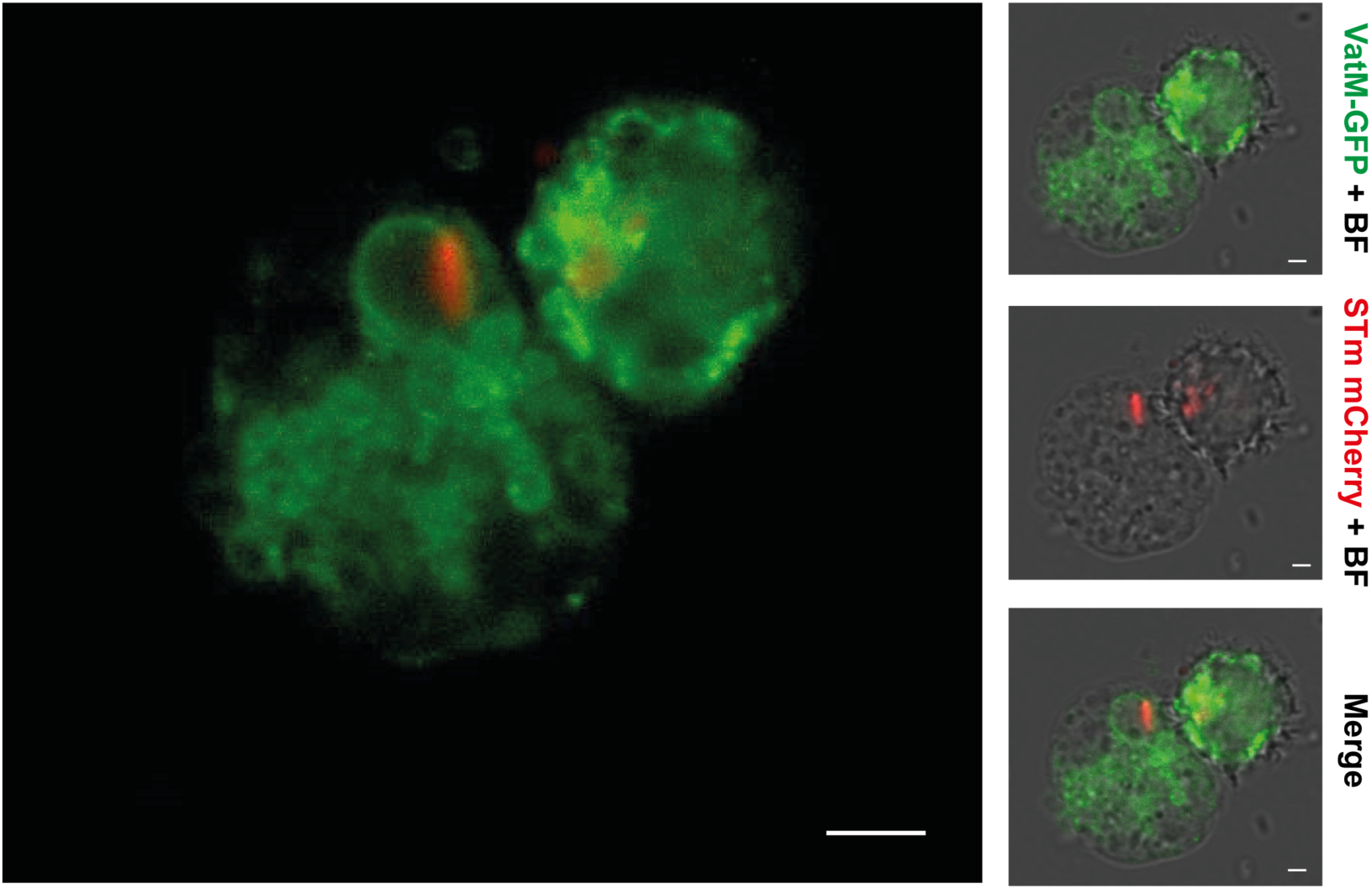
*S.* Typhimurium resides in a spacious vacuolar compartment surrounded by the vacuolar ATPase. *S.* Typhimurium 14028s constitutively expressing mCherry was used to infect the axenic strain *D. discoideum* AX2 VatM-GFP. Images were acquired at 4.5 h post infection with a Zeiss LSM 710 confocal microscope using ZEN 2012 Black software (Zeiss), and edited using FIJI software. The green fluorescence corresponds to the fusion protein VatM-GFP and the red fluorescence corresponds to bacteria expressing mCherry. BF: brightfield. Bar is 2 μm.

### Preparation of intact SCVs from *D. discoideum* infected with *S*. Typhimurium

We pursued to characterize the protein content of these *Salmonella-*containing vacuolar compartments in *D*. discoideum. Therefore, we adapted a protocol originally developed to obtain a subcellular fraction highly enriched in intact SCVs from infected HeLa cells (43). As described in the **Supplementary Figure S2**, infected and control amoebae were lysed, and the post-nuclear supernatants obtained were loaded on top of linear 10-25% Optiprep gradients. After centrifugation, 12 fractions obtained from each sample were analyzed. As shown in **Figure 2**, in control samples bacteria accumulated in fractions F8 and F9 (1.12-1.13 g/cm^3^), while in samples from infected amoebae we noted a shift in this distribution with bacteria also accumulating at the lower-density fractions F6 and F7 (1.09-1.10 g/cm^3^). The presence of this differential bacterial distribution between infected and control samples is consistent with the previously obtained data of bacteria residing inside vacuoles in infected human cells (43). To confirm the presence of vacuoles containing bacteria in our preparations, we performed an anti-*Salmonella* ELISA-based assay (43). In this assay, an immobilized anti-*Salmonella* antibody was used to capture free bacteria present in each fraction that were subsequently quantified using a biotin-conjugated anti-*Salmonella* antibody. The samples were then analyzed before and after an osmotic shock treatment to break the SCV and release all bacteria residing in this compartment. This procedure provided information on the respective fractions of vacuolar-bound *Salmonella* and free *Salmonella*. As shown in **Supplementary Figure S3**, the amount of *Salmonella* detected by our assay in fractions F6 to F8 from infected cells increased after the osmotic shock, while there was no increase in the number of *Salmonella* detected in the lower density fraction before and after the osmotic shock treatment in the case of control samples. Together, these results indicate that fractions F6 and F7 were enriched in intact vacuoles containing *Salmonella*. Consequently, we used this fractionation procedure to analyze the protein composition of the SCV.

**Figure 2.**
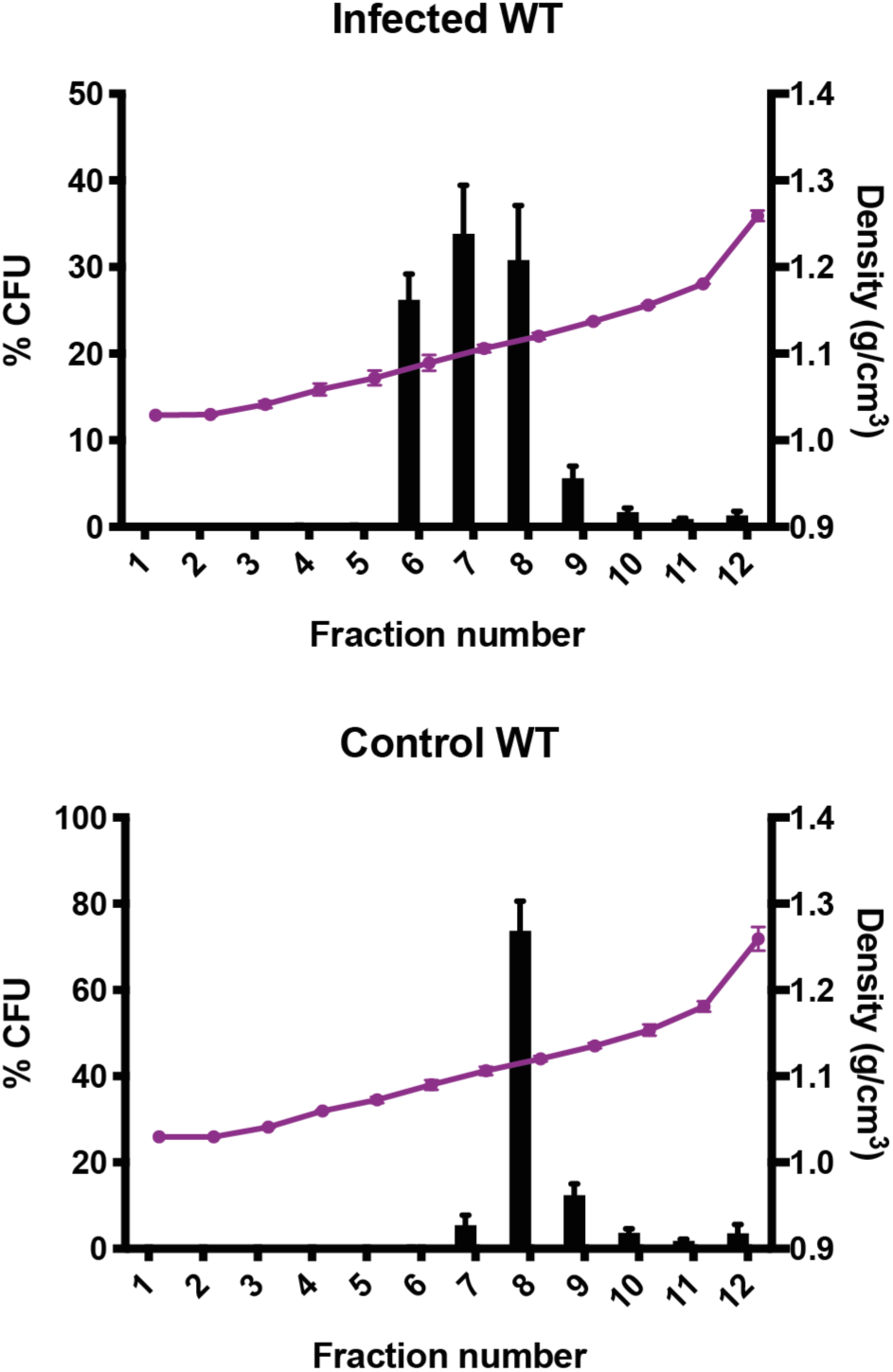
Subcellular fractionation of *D. discoideum* cells infected with *S*. Typhimurium WT and uninfected control cells. Graphs show the CFU distribution and density per fraction of PNS samples obtained from cells infected with *S*. Typhimurium 14028s (upper panel) or PNS from uninfected control cells spiked with a known amount of *S*. Typhimurium 14028s (lower panel). Each graph shows mean values ± SD from three independent experiments.

Additionally, we wanted to evaluate the impact of the absence of effectors SopB and SifA on the proteome of the SCV in *D. discoideum*. To this end, we performed our infection assays and fractionation experiments using Δ*sopB* and Δ*sifA* mutants obtained from our wild-type *S*. Typhimurium strain. In the case of samples from amoebae infected with the Δ*sopB* and Δ*sifA* mutants, we also observed the shift in bacterial distribution between control and infected samples (**Figure 3**), although the increase of bacteria in F6 after osmotic shock was not as evident as in samples from amoebae infected with the wild-type strain (**Figure 2**). When these samples were analyzed by our anti-*Salmonella* ELISA, we observed an increase in *Salmonella* detected after the osmotic rupture of the vacuoles (**Supplementary Figure S4**), indicating the presence of intact vacuoles containing the *Salmonella* strains in the samples obtained from infected cells. For each bacterial strain infecting *D. discoideum*, four biological replicates were obtained for proteomic analyses.

**Figure 3.**
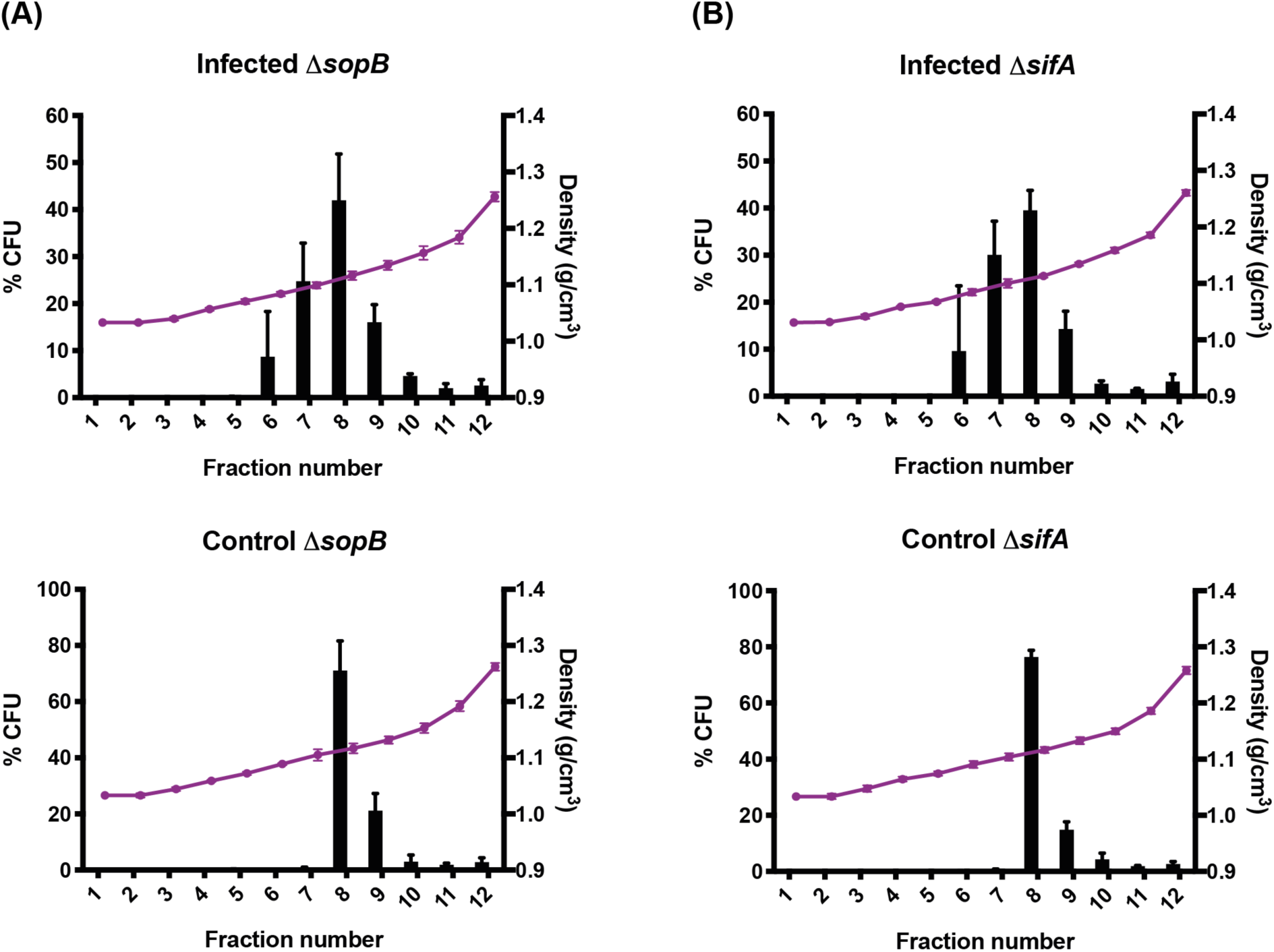
Subcellular fractionation of *D. discoideum* cells infected with *S*. Typhimurium Δ*sopB* and Δ*sifA* mutants and uninfected control cells. Graphs show the CFU distribution and density per fraction of PNS samples obtained from cells infected with *S*. Typhimurium Δ*sopB* **(A, upper panel)** or Δ*sifA* **(B**, **upper panel)**, or uninfected control cells spiked with a known amount of *S*. Typhimurium Δ*sopB* **(A, lower panel)** or Δ*sifA* **(B, lower panel)**. Each graph shows mean values ± SD from four independent experiments.

### Proteomic analysis of SCVs recovered from infected *D. discoideum*

Fractions F6 and F7 from each experiment were selected for proteomic analysis. These fractions were subjected to protein precipitation, and the proteins from both fractions per experiment were pooled and analyzed by LC-MS/MS. Proteomic data analysis was performed using the PatternLab for Proteomics 4.0 software (44). We first analyzed the data obtained from amoebae infected with the wild-type strain (WT-infected samples) and the corresponding uninfected control samples. For WT-infected samples 1,319 proteins were identified, while for control samples 1,389 proteins were identified. After manual curation of these two datasets to remove proteins corresponding to contaminants and the few bacterial proteins detected, we identified 714 proteins that were shared between both conditions, 91 proteins present only in WT-infected samples, and 137 proteins present only in control samples (**Figure 4A**). Among the proteins present only in WT-infected samples we found proteins related to intracellular trafficking; proteins involved in multivesicular body formation; motor proteins and actin related proteins. We also found proteins involved in degradative pathways such as E2 and E3 ubiquitin ligases; and different subunits of the COP9 signalosome. The complete list of proteins presents exclusively enriched in WT-infected samples is presented in **Supplementary Table S1**. Next, we analyzed the proteins found both in WT-infected and control samples to pinpoint those that were enriched in samples obtained from infected cells, according to their normalized spectral abundance factors (NSAF). To do this, we used the PatternLab TFold module to generate a volcano plot according to the fold change (F-change) and P-value of each protein (**Figure 4B**). Light blue dots (NC in **Supplementary Table S2**) represent proteins that do not satisfy neither F-change nor statistic criteria, and thus were considered as unchanged between the two conditions. Teal dots (NS in **Supplementary Table S2**) satisfy the F-change criterion, but not the statistical one. Light purple dots (C in **Supplementary Table S2**) correspond to low abundant proteins satisfying both the F-change and Q-value criteria, but due to the low number of spectra they deserve further validation. Finally, dark purple dots (SC in **Supplementary Table S2**) correspond to proteins satisfying all statistical filters and represent the over- and under-represented proteins between infected and control samples.

**Figure 4.**
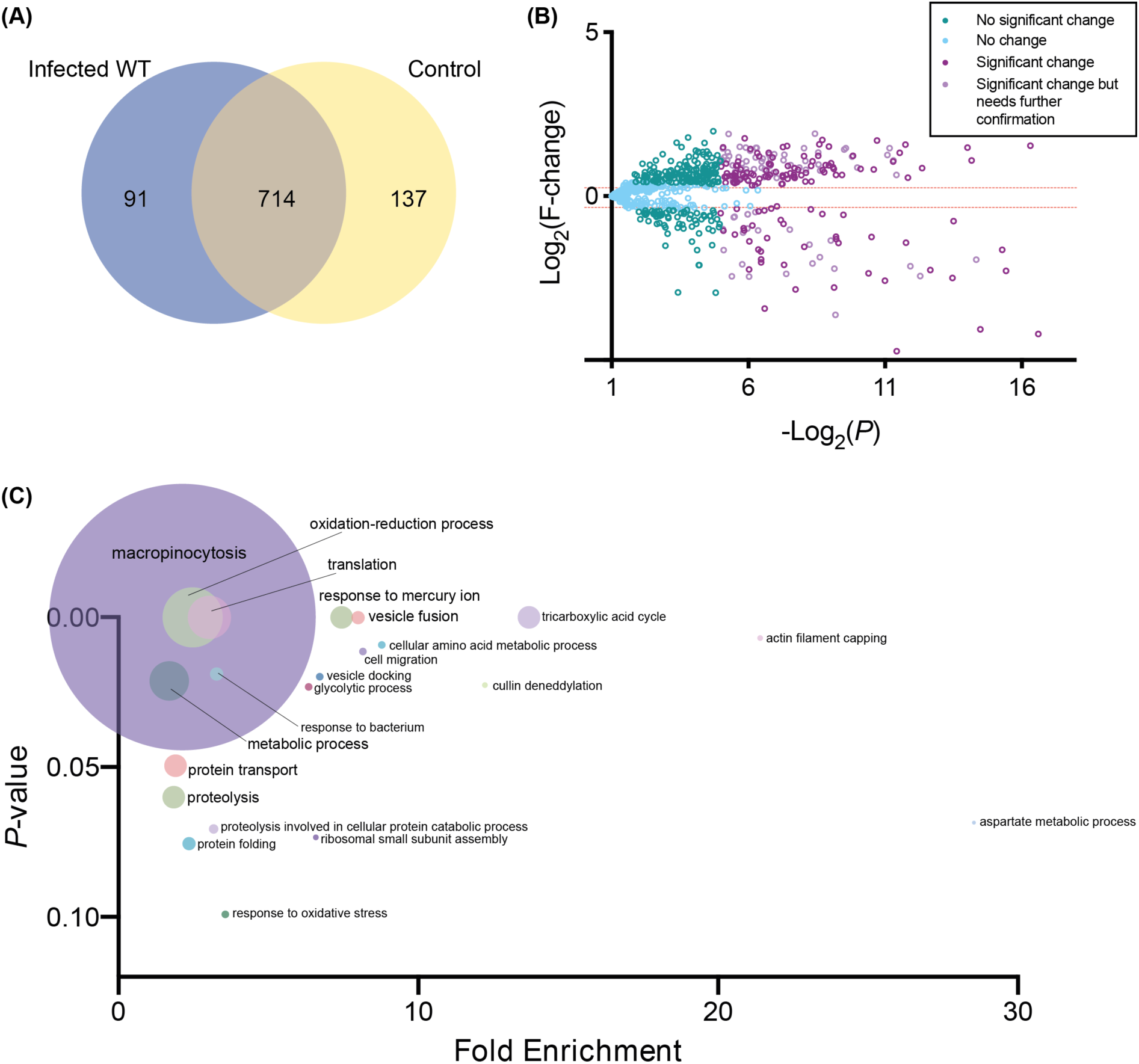
Proteomic analysis of WT-infected and uninfected control samples. **(A)** Venn diagram showing the distribution of proteins identified in samples from cells infected with the WT strain or uninfected control samples. **(B)** Volcano plot generated using the PatternLab for Proteomics TFold module (FDR of 0.05, F-stringency of 0.03 and Q-value of 0.05). For this particular dataset, when a protein presented a F-change > 1.25 and a *P*-value < 0.05 it was considered to be enriched, and when it presented a F-change < -1.24 and a *P*-value < 0.05 it was defined as underrepresented in WT-infected samples. Each dot represents a protein identified in 3 replicates of all conditions, plotted according to its *P*-value (Log_2_(*P*)) and fold change (Log_2_(F-change)). **(C)** Gene ontology functional clustering analysis using DAVID, showing the biological processes enriched in infected samples. The size of the dot represents the number of proteins enriched in that biological process. All data on these figures can be found in **Supplementary Tables S1** and **S2**.

We determined the number of proteins specifically enriched in WT-infected samples: 90 proteins in the SC category and 62 in the C category. Among them, we found trafficking related proteins, GTPases (Rab5B and RacE) and actin-related proteins. Again, we found proteins related to degradative compartments such as cathepsin D, LmpB (lysosome membrane protein 2-B) and the autophagy marker protein Atg8 (also known as LC3). The complete list of proteins differentially represented between WT-infected and control samples is presented in **Supplementary Table S2**.

Next, we analyzed the data from samples obtained from amoebae infected with either Δ*sopB* or Δ*sifA* mutants using the same parameters employed for the analysis of the WT-infected samples. In the case of the Δ*sopB*-infected samples 1,520 proteins were identified, while 1,414 proteins were identified in the corresponding control samples. After manual curation of these two datasets we identified 779 proteins that were shared between both experimental conditions, 253 proteins present only in Δ*sopB*-infected samples, and 46 proteins present only in control samples (**Figure 5A**). In the case of the Δ*sifA*-infected samples 1,542 proteins were identified, while 1,423 proteins were identified in the corresponding control samples. By comparing these two datasets after manual curation we identified 746 proteins shared between both experimental conditions, 228 proteins present only in Δ*sifA*-infected samples, and 47 proteins present only in control samples (**Figure 6A**).

**Figure 5.**
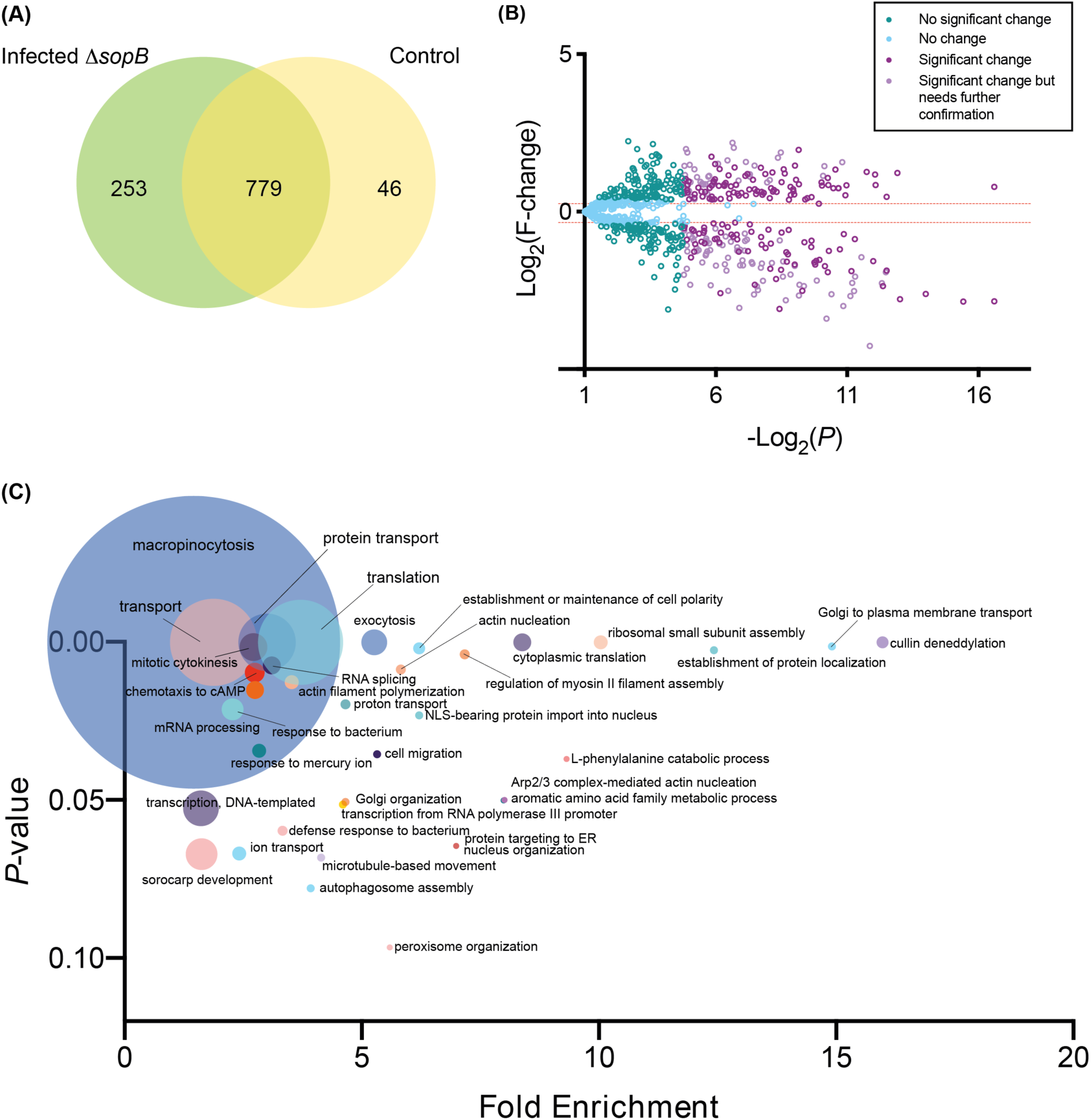
Proteomic analysis of Δ*sopB*-infected and uninfected control samples. **(A)** Venn diagram showing the distribution of proteins identified in samples from cells infected with the Δ*sopB* mutant or uninfected control samples. **(B)** Volcano plot generated using the PatternLab for Proteomics TFold module (FDR of 0.05, F-stringency of 0.03 and Q-value of 0.05). For this comparison, when a protein presented an F-change > 1.19 and a *P*-value < 0.05 it was considered to be enriched and when it presented an F-change < -1.27 and a *P*-value < 0.05 it was defined as underrepresented in Δ*sopB-*infected samples. Each dot represents a protein identified in 4 replicates of all conditions, plotted according to its *P*-value (Log_2_(*P*)) and fold change (Log_2_(F-change)). **(C)** Gene ontology functional clustering analysis using DAVID, showing the biological processes enriched in infected samples. The size of the dot represents the number of proteins enriched in that biological process. All data on these figures can be found in **Supplementary Tables S3** and **S4**.

**Figure 6.**
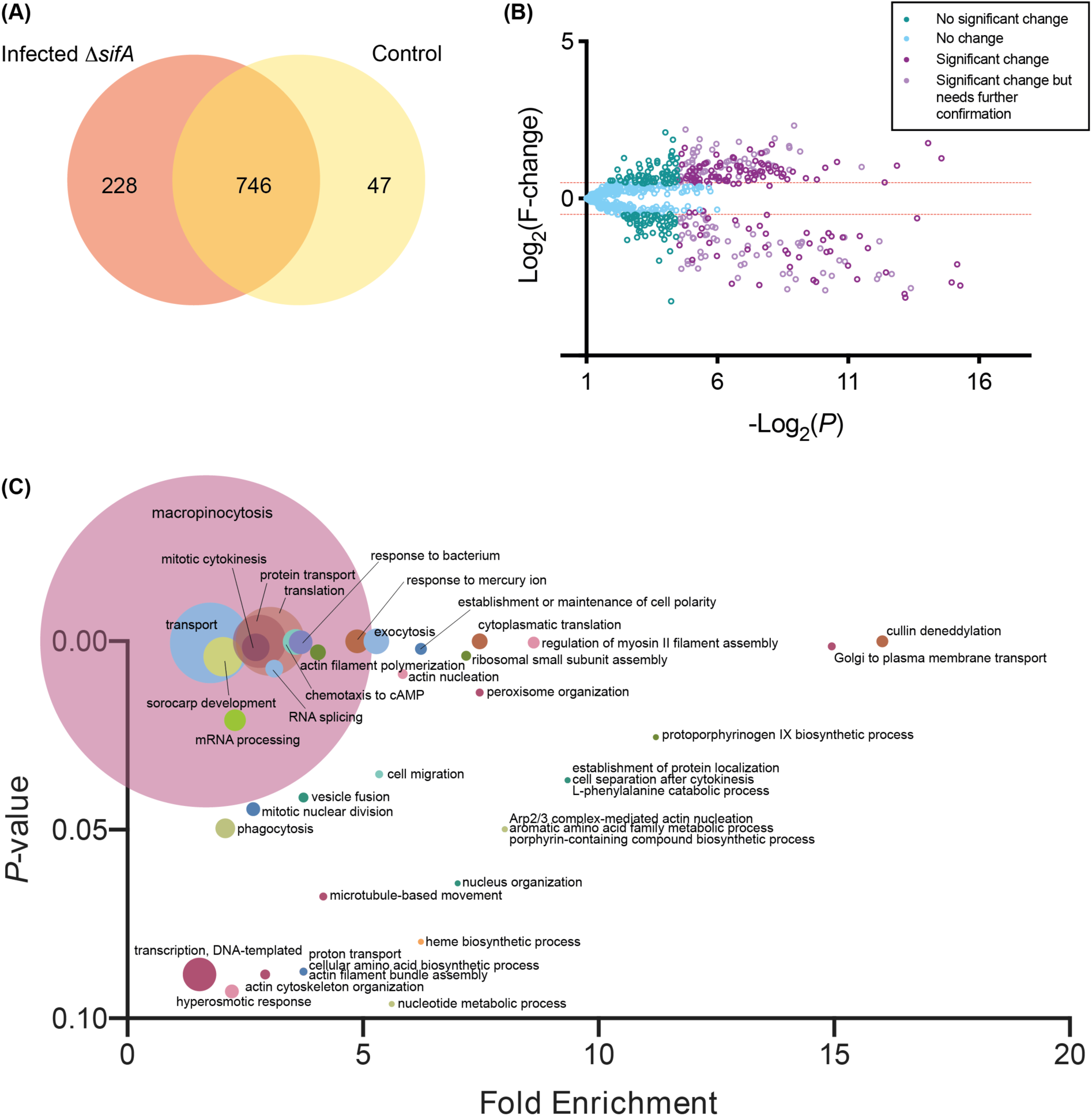
Proteomic analysis of Δ*sifA*-infected and uninfected control samples. **(A)** Venn Diagram showing the distribution of proteins identified in samples from cells infected with the Δ*sifA* mutant or uninfected control samples. **(B)** Volcano plot generated using the PatternLab for Proteomics TFold module (FDR of 0.05, F-stringency of 0.04 and Q-value of 0.05). For this dataset, when a protein presented an F-change > 1.32 and a *P*-value < 0.05 it was considered to be enriched, and when it presented an F-change < -1.32 and a *P*-value < 0.05 it was considered as underrepresented in Δ*sifA*-infected samples. Each dot represents a protein identified in 4 replicates of all conditions, plotted according to its *P*-value (Log_2_(*P*)) and fold change (Log_2_(F-change)). **(C)** Gene ontology functional clustering analysis using DAVID, showing the biological processes enriched in infected samples. The size of the dot represents the number of proteins enriched in that biological process. All data on these figures can be found in **Supplementary Tables S5** and **S6**.

Among the proteins detected exclusively in Δ*sopB*-infected samples, we found proteins related to intracellular trafficking, components of the Exocyst complex; Ras guanine nucleotide exchange factors (GEFs) and Rho GTPase-activating proteins (GAPs); motor proteins; Ubiquitin related enzymes; and autophagy related proteins Atg3 and Atg12. The complete list of proteins exclusively present in Δ*sopB*-infected samples is presented in **Supplementary Table S3**. After using the TFold module, we determined that 95 proteins in the SC category and 40 in the C category were specifically enriched in Δ*sopB*-infected samples (**Figure 6B**). Among the proteins specifically enriched in Δ*sopB*-infected samples, we found those related to trafficking (Syntaxin 7, Vti1A, Vps35, ExoC7, GefH); small GTPases (RacC, RacE, RapA and Rab14) and actin-related proteins. Noteworthy, several Rab GTPases were underrepresented in Δ*sopB*-infected samples, such as Rab1, Rab2, Rab5, Rab6 and Rab8. The complete list of proteins differentially represented between Δ*sopB*-infected and control samples is presented in **Supplementary Table S4**.

Among the proteins present exclusively in Δ*sifA*-infected samples, we found the autophagy related proteins Atg5 and Atg3; the vacuolar proteins Vps37 and Vps51; GAPs and GEFs. The complete list of proteins that are present exclusively in Δ*sifA-*infected samples is presented in **Supplementary Table S5**. Using the TFold module to analyze the distribution between proteins in these two conditions we determined that 88 proteins in the SC category and 63 in the C category were specifically enriched in Δ*sifA*-infected samples (**Figure 7B**). Of note, we found proteins related to the same pathways to those present exclusively in Δ*sopB*-infected samples, such as trafficking proteins, the small GTPases RapA and RacE; the guanine nucleotide binding protein GpbB; GefH; and actin-related proteins. As in the case of the Δ*sopB*-infected samples, several Rab GTPases were found to be underrepresented in Δ*sifA*-infected samples, including Rab1, Rab2, Rab5 and Rab8. The complete list of proteins differentially represented in Δ*sifA*-infected and control samples is presented in Supplementary Table S6.

**Figure 7.**
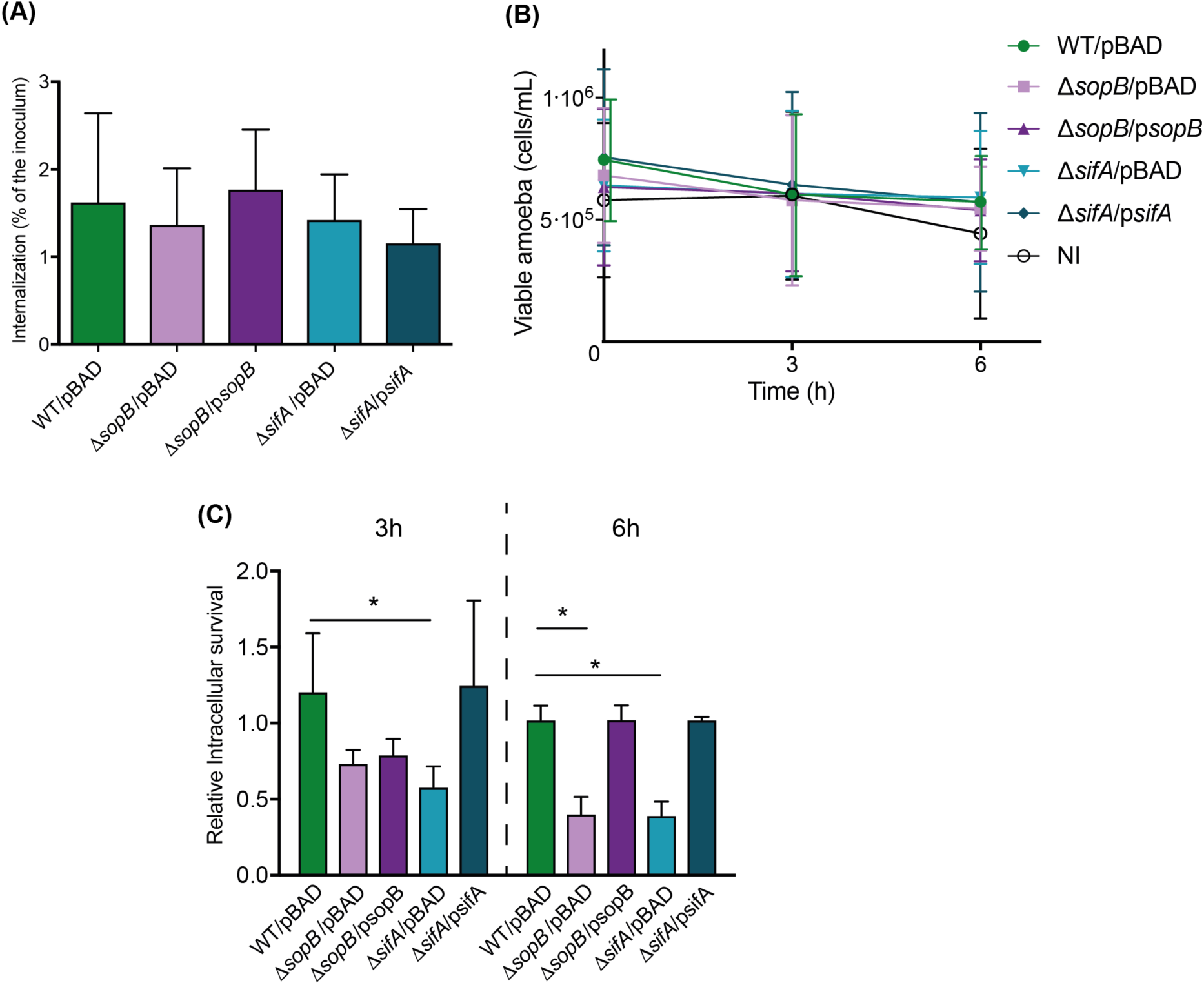
Intracellular survival of WT, Δ*sopB* and Δ*sifA* strains of *S*. Typhimurium in *D. discoideum*. **(A)** Internalization expressed as the percentage of intracellular bacteria at t=0 relative to the initial inoculum. Statistical significance was determined using a one-way ANOVA. **(B)** Population of viable amoebae at each time post infection. Statistical significance was determined using a one-way ANOVA. **(C)** Intracellular survival expressed as CFU/cell at t=3 h or t=6 h divided by the CFU/cell at t=0. Statistical significance was determined using a two-way ANOVA with Dunnet’s test (* = *P* < 0.05). All graphs show mean values ± SEM of at least three independent assays.

**Figure 8.**
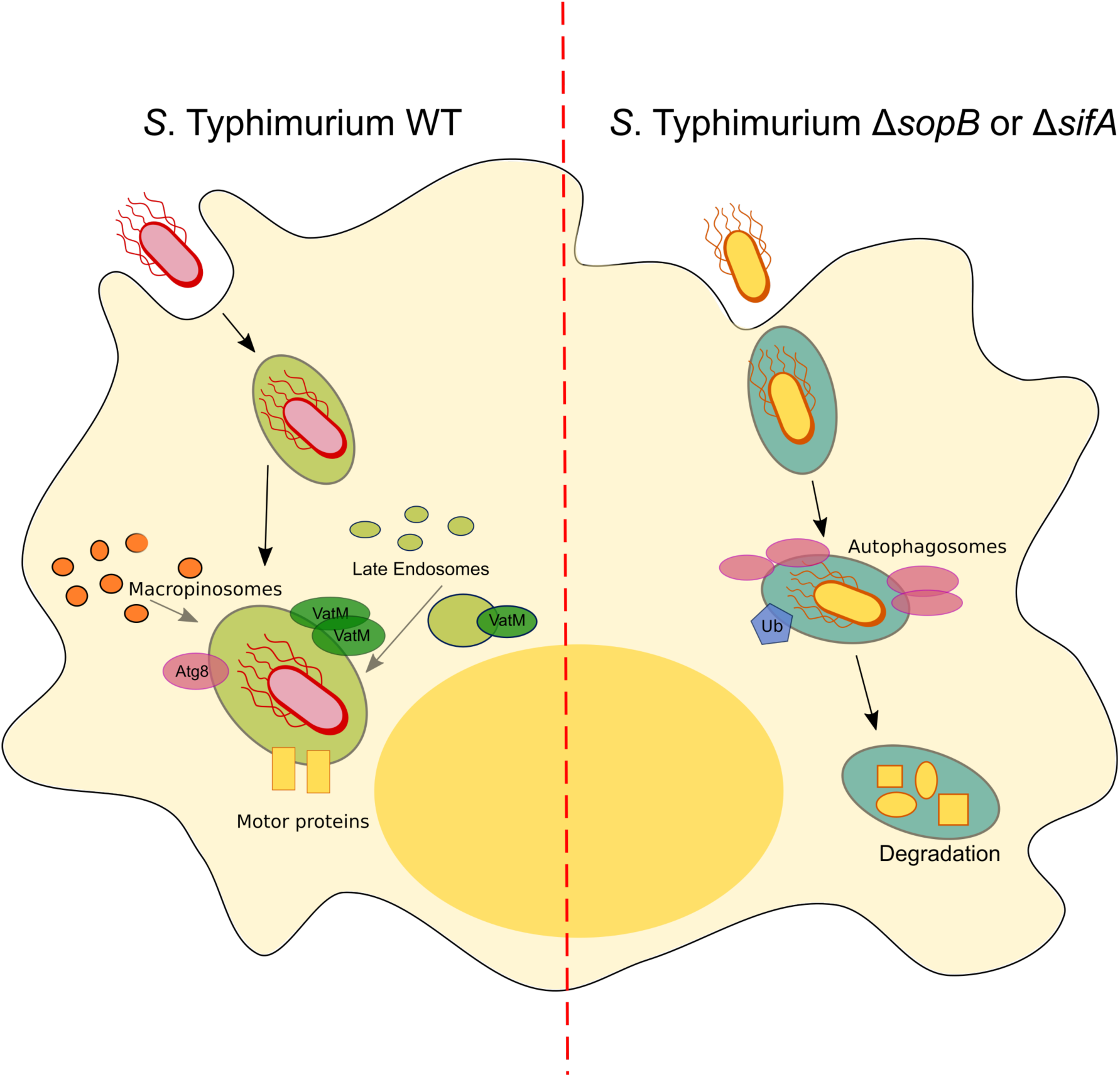
Model for the biogenesis of the SCV in *Dictyostelium discoideum*. After internalization of *Salmonella* by *D. discoideum*, the pathogen resides in a specialized vacuolar compartment that avoid phagosome degradation by exploiting the activity of effectors SopB and SifA. These proteins allow the recruitment of the autophagy machinery and other intracellular membrane-associated proteins to the SCV in order to sustain the intracellular survival of *Salmonella* in this organism. The absence of effectors SopB and/or SifA results in the generation of a vacuolar compartment destined for degradation.

In addition to the analyses described above, we performed a functional clustering analysis using DAVID (Database for Annotation, Visualization and Integrated Discovery) (45, 46) for all the proteins identified in the different experimental conditions and those enriched in samples infected with different *Salmonella* strains. The overview of the biological processes associated with proteins detected in each experimental condition is shown in **Figures 5C**, **6C** and **7C**. In all cases, most proteins appeared as groups related to processes of intracellular trafficking, such as macropinocytosis, exocytosis, or motor protein assembly.

Overall, our fractionation procedure followed by quantitative proteomics allowed us to characterize the SCV composition during *D. discoideum* infections with *S*. Typhimurium. Furthermore, differences in SCV proteomes from amoebae infected with WT, Δ*sopB* and Δ*sifA* strains suggest that the effectors SopB and SifA are required to generate a mature vacuolar compartment that escapes from degradative pathways and sustains the intracellular lifestyle of the pathogen in this host. For instance, the presence of autophagy proteins Atg8 and Atg12 in SCVs from amoebae infected with the WT strain highlights the importance of autophagy to support *Salmonella* survival in this organism. In contrast, the presence of a different set of autophagy markers (including Atg3, Atg5 and Atg12), together with an increased number of proteins associated with lysosomal degradation in SCVs from amoebae infected with Δ*sopB* and Δ*sifA* mutant strains indicate that these mutants reside in a vacuolar compartment destined for degradation.

### SopB and SifA effectors are required for intracellular survival of *S*. Typhimurium in *D. discoideum*

Considering the results from our proteomic analyses, we decided to evaluate the intracellular survival of Δ*sifA* and Δ*sopB* mutants in *D. discoideum*. We anticipated that these mutants would be contained in a vacuolar compartment unable to support the intracellular survival of the pathogen. Therefore, we performed infection assays in *D. discoideum* under conditions that have been described by our group (27, 28) (**Figure 7**). We found that the number of internalized bacteria was similar between the different strains (**Figure 7A**), indicating that the deletion of genes *sopB* and *sifA* does not affect the uptake of *S*. Typhimurium by *D. discoideum*. Then, we evaluated the intracellular survival of these strains and observed that the Δ*sifA* mutant presented defects at 3 h post infection, and that this phenotype became more prominent at 6 h post infection (**Figure 7B**). Similarly, the Δ*sopB* mutant also presented growth defects at 6 h post infection (**Figure 7B**). The phenotype shown by each mutant was reverted by the presence of a derivative of plasmid pBAD-TOPO harboring a wild-type copy (including the promoter region) of the corresponding gene. No strain significantly changed the number of viable amoebae during the course of the experiment (**Figure 7C**), indicating that the differences observed in the titers of intracellular Δ*sopB* and Δ*sifA* mutants are not attributable to changes in the number of viable amoebae.

Thus, our results indicate that SopB and SifA effectors are required for the intracellular survival of *S*. Typhimurium in *D. discoideum*. The survival defect shown by Δ*sopB* and Δ*sifA* mutants correlate with the differential recruitment of host degradation factors observed in our proteomics analysis.

## Discussion

### *S*. Typhimurium resides in a vacuolar compartment in *D. discoideum*

In the present study, we first analyzed the presence of a vacuolar compartment that contained *S*. Typhimurium in infected *D. discoideum* cells. To evaluate the presence of such compartment, we employed confocal microscopy and performed bacterial infections using a *D. discoideum* reporter strain that expresses a component of the vATPase called VatM fused to GFP. This enzyme uses ATP hydrolysis to transport protons across membranes and are composed of two subcomplexes: V_1_ and V_0_ (40). VatM is part of the V_0_ complex and has been localized to the membranes of the contractile vacuole in *D. discoideum* (47); and in membranes of the endolysosomal system (48), in which vATPase acidifies the lumen of endosomes. It is also important to note that vATPase is recruited to phagosomal and endolysosomal membranes of other cellular models, such as epithelial cells and macrophages. Furthermore, vATPase is also present in the SCV in HeLa and macrophage cells (33, 41, 49, 50). Therefore, we tested the presence of the vATPase in *D. discoideum* VatM-GFP infected with *S*. Typhimurium constitutively expressing mCherry. In our experiments, we observed the presence of the VatM-GFP marker on vacuoles containing *S*. Typhimurium at 3 and 4.5 h post infection (**Figure 2** and **Supplementary Figure S1**). This confirmed that *S*. Typhimurium is contained in a vacuolar compartment in *D. discoideum*, as described by other groups (25), and that this compartment is a VatM^+^ vacuole. Other pathogens such as *Mycobacterium marinum* and *Legionella pneumophila* also reside within a membrane-bound compartment in *D. discoideum* (21, 51), each of them with their particular characteristics. Of note, in the case of *M. marinum* there is a selective exclusion of the vATPase from the *Mycobacterium*-containing vacuole (MCV) to avoid acidification of the vacuole (52) and in the case of *L. pneumophila*, VatA (another vATPase subunit) is also excluded from the *Legionella*-containing vacuole (LCV) (53).

### The proteome of the vacuolar compartment containing *S*. Typhimurium in *D. discoideum*

The SCV has been predominantly characterized in infected HeLa cells, and the proteome of the compartment in this cell line has been reported (43). This study identified about 400 proteins, and the data showed a significant enrichment in proteins from several organelles, including ER, early and late endosomes trans-Golgi network and lysosomes, along with vesicle-transport related proteins and cytoskeleton proteins that may act stabilizing SIFs. It also underlined the importance of the ER and ER-contacting points in defining the fate of intracellular *S*. Typhimurium in HeLa cells, which are used as a model for epithelial cells mainly because of their easy manipulation for cell biology studies. *Salmonella* replication in these cells is highly permissive, in contrast to replication in macrophages and other phagocytic cells. For this reason, our analysis of the compartment containing *Salmonella* in *D. discoideum* could be more comparable to studies performed in macrophages and to other bacteria-containing compartments in phagocytic cells. Only recently, the isolation of SCVs from human THP-1 macrophages using paramagnetic nanoparticles attached to the surface of *S*. Typhimurium has been reported (54). The study performed an initial characterization of classical SCV-associated proteins, such as Rab5 and LAMP-1, but did not analyze the proteome of this vesicular compartment. Furthermore, the nanoparticles attached to the bacterial surface may influence the internalization pathway, which alters the initial compartment formed and influences how *S*. Typhimurium subverts the host trafficking pathways (55).

In a related study in *L. pneumophila*, the proteomes of LCVs were compared from *D. discoideum* and RAW264.7 macrophages (56). This highlighted the similarities between both compartments in the two host cells. In this study, numerous LCV proteins identified from infected RAW264.7 macrophages have been described in the literature, including several small GTPases (57). In addition, novel LCV components were identified, such as the small GTPases Rab2A, Rab6, Rab11A, Rab18, Rab32A, RacB, RacE and RapA; some GTPase modulators (Rab, Ras and Rho GAPs); SNARE proteins; and Ser/Thr kinases (56). When compared to proteins identified in LCVs from *D. discoideum*, the authors found a considerable number of proteins implicated in the same signaling pathways, including Ser/Thr and Tyr protein kinases and phosphatases, cyclin-dependent kinases, small GTPases of the Rho/Rac, Ras or Ran families, GTPase modulators, ubiquitin-dependent factors, multivesicular body (MVB) proteins, cargo receptors, dynamin-like GTPases, sorting nexins (SNXs), syntaxins, motor proteins (dynein, kinesin and myosin), and factors implicated in microtubule dynamics (56). Interestingly, our proteome results are similar regarding the type of SCV proteins identified and the biological processes involved, as we found SNARE proteins, motor proteins, ubiquitin ligases, Ser/Thr kinases and phosphatases, and MVB proteins, among others. In contrast, our results differ particularly in the Rab GTPases that are enriched in the LCV proteome, as we found that these proteins are not enriched in SCV proteomes. These differences are likely due to the activity of specific effectors secreted by the different bacteria that generate distinct pathogen-specific niches.

In the SCV proteomes obtained from *D. discoideum* infected with the mutant strains we found several proteins involved in degradative pathways, such as ubiquitin ligases, COP9 signalosome (a type of proteasome that cleaves ubiquitin conjugates and ubiquitin-like protein conjugates, among other targets) (58, 59) and autophagy related proteins. Autophagy is a highly-conserved process from yeast to mammals, and many genes associated with autophagy (*atg*) are conserved in amoebae, plants, worms and mammals, emphasizing the importance of this process. Autophagy also controls infections caused by intracellular pathogens, as they can be captured in autophagosomes for degradation. In *D. discoideum*, autophagy is the main process that allows this organism to fight intracellular pathogens that escape the endolysosomal degradation pathway (60, 61). Pathogens such as *M. marinum* have been described to subvert autophagy in *D. discoideum* by inducing the autophagy pathway via transient inhibition of TORC1 activity at early stages of interaction, and avoiding being killed inside autolysosomes by blocking the autophagic flux (21). This results in the accumulation of membranes and cytoplasmic material in the MCV, which might support bacterial survival within this niche.

Similar mechanisms have been described in *Salmonella*, as ruptured SCVs are recognized by galectins (cytoplasmic lectins that bind specific carbohydrate modifications within the ruptured SCV), which subsequently recruit adaptors and autophagosomes (62). Moreover, several reports indicate that autophagy targets cytoplasmic *Salmonella* for degradation (62–64). However, other studies demonstrated a role of the autophagy machinery in the repair of damaged SCV membranes caused by T3SS_SPI-1_ activity (65), and that the autophagy machinery associated with cytosolic *Salmonella* and promotes intracellular replication (66). Current studies by our group show that *S*. Typhimurium subverts the autophagy machinery in both *D. discoideum* and RAW264.7 macrophages by means of effector proteins secreted by T3SS_SPI-1_ (Urrutia et al, manuscript in preparation). Yet, another study determined that autophagy is necessary to avoid intracellular replication of *S*. Typhimurium in *D. discoideum*, as amoebae carrying null mutations in genes linked to the autophagy pathway infected with *S*. Typhimurium show a decrease in lifespan and an increased bacterial intracellular replication (67). Taking these studies together, our proteomic analysis highlights the importance of the autophagic machinery in the fate of the vacuolar compartment involved in *S*. Typhimurium survival in *D. discoideum*.

### *S*. Typhimurium requires effectors SopB and SifA to survive intracellularly in *D. discoideum*

The role of the effector proteins SopB and SifA in the intracellular survival of *Salmonella* in other hosts has been widely reported. *In vitro*, it has been shown that SopB contributes to invasion of HeLa cells (32), but is dispensable for intracellular replication in intestinal Henle-407 cells (31). On the other hand, this effector it is required for intracellular growth of *Salmonella* in bone marrow-derived macrophages (31), which is similar to the phenotype we observe in *D. discoideum*. SopB also plays a critical role in the size control of the SCV and its stability (68). In the case of SifA, it has been shown that *sifA* mutants are strongly attenuated in mice (69), but they show a higher percentage of escape from the SCV and hyperreplicate in the cytosol of HeLa cells (70). In addition, *sifA* mutants are defective for intracellular replication in restrictive cell lines such as Swiss 3T3 fibroblasts and RAW 264.7 macrophages (69). Hyperreplication of *Salmonella* has not been described in *D. discoideum* and we have no evidence of this particular phenotype in this model. Moreover, it has been shown that *Salmonella* requires SopB and SifA to survive and grow intracellularly in mammalian phagocytic cells (31, 69). Consistently with these phenotypes, our results show that both effectors play a critical role in the intracellular survival of *S*. Typhimurium in *D. discoideum*.

From our work, we propose a model in which *Salmonella* is internalized by *D. discoideum* and resides in a specialized vacuolar compartment to avoid phagosome degradation by exploiting the activity of specific effectors secreted through T3SS_SPI-1_ and T3SS_SPI-2_. For instance, this specialized vacuole can be modified by the action of effector proteins SopB and SifA, allowing the recruitment of the autophagy machinery (65, 66) and other intracellular membrane-associated proteins allowing the survival of *S*. Typhimurium within *D. discoideum*, and its replication at later times of infection (28). Altogether, our results indicate that the SCV in *D. discoideum* presents similarities with other pathogen-containing vacuoles, both in this model host and in other phagocytic cells such as macrophages. This highlights the importance of *D. discoideum* as a model to study *Salmonella* survival in phagocytic amoebae. In the future, the data generated in this work can be further explored as a starting point to determine other cellular and bacterial proteins involved in biological processes related to the interaction of *S*. Typhimurium with *D. discoideum*.

## Materials and Methods

### Bacterial strains and growth conditions

The bacterial strains used in the present study are listed in **Supplementary Table S7**. All *S*. Typhimurium strains are derivatives of the wild-type virulent strain 14028s (71). Bacteria were routinely grown in Luria-Bertani (LB) medium (10 g/L tryptone, 5 g/L yeast extract, 5 g/L NaCl) at 37°C with aeration. LB medium was supplemented with ampicillin (Amp; 100 mg/L) or kanamycin (Kan; 75 mg/L) as appropriate. Media were solidified by the addition of agar (15 g/L). All procedures involving the use of pathogenic organisms were conducted following the guidelines in the Biosafety Manual of the National Commission of Scientific and Technological Research (CONICYT), and were approved by the Institutional Biosafety Committee of Universidad de Chile, Campus Norte.

### Construction of mutant strains, cloning and complementation

*S*. Typhimurium mutants with specific deletions of *sopB* and *sifA* genes and the concomitant insertion of a Kan-resistance cassette were constructed using the Lambda Red recombination method (72) with modifications (73). PCR amplification of the resistance cassette present in plasmid pCLF4 (GenBank accession number HM047089) was carried out under standard conditions using primers listed in **Supplementary Table S8**. *S*. Typhimurium strain 14028s carrying plasmid pKD46, which encodes the Red recombinase system, was grown to an OD_600nm_ of 0.5-0.6 at 30°C in LB medium containing Amp and L-arabinose (10 mM). Bacteria were made electrocompetent by sequential washes with ice-cold sterile 10% glycerol, and transformed with ∼500 ng of each purified PCR product. Transformants were selected at 37°C on LB agar containing Kan. The presence of each mutation was confirmed by PCR amplification using primers flanking the sites of substitution (**Supplementary Table S8**). Finally, each mutation was transferred to the wild-type background by generalized transduction using phage P22 HT105/1 *int*-201, as described (74).

For complementation assays, genes *sopB* and *sifA* were PCR amplified from DNA obtained from the wild-type strain using primers flanking the promoter region and ORF of each gene (**Supplementary Table S8**). PCR products were ligated to the pBAD-TOPO vector (Invitrogen) and transformed into chemically competent *E. coli* TOP10 cells (Invitrogen), according to the manufacturer instructions. Transformants were selected on LB agar containing Amp. Recombinant plasmids with either gene cloned in the same orientation as the *P_araBAD_* promoter were identified by PCR using combinations of primers listed in **Supplementary Table S8**. Each plasmid was purified using the QIAPrep Spin Miniprep Kit (Qiagen) and transformed in the corresponding mutant strain for complementation assays. In addition, the wild-type and mutant strains containing the empty pBAD-TOPO vector were also generated as controls.

### *Dictyostelium* strains and growing conditions

*D. discoideum* strains AX2 (DBS0235519) (75) and AX2 VatM-GFP (DBS0235537) (40) were obtained from Dicty Stock Center (76–78), and cultured according to standard protocols (79). Amoebae were maintained at 23°C in SM agar (10 g/L tryptone, 1 g/L yeast extract, 1.08 g/L MgSO_4_ x 7H_2_O, 1.9 g/L KH_2_PO_4_, 0.78 g/L K_2_HPO_4_ x 3H_2_O, 10 g/L glucose, 20 g/L agar agar) growing on top of a confluent lawn of *Klebsiella aerogenes* DBS0305928 until phagocytosis plaques were visible. Growing cells were transferred to liquid HL5 medium (14 g/L tryptone, 7 g/L yeast extract, 0.35 g/L Na_2_HPO_4_, 1.2 g/L KH_2_PO_4_, 15.2 g/L glucose) containing Amp (100 µg/mL) and streptomycin (Str; 300 µg/mL), and incubated at 23°C in tissue culture flasks when adherent cells were needed, or in glass flasks with agitation (180 rpm) when cells in suspension were needed. Cells were subcultured and used in the different assays when they reached 70-80% confluence in tissue culture flasks or when they reached exponential phase (1-2 x 10^6^ cells/mL) when cultured in suspension. HL5 medium was supplemented with G418 (10 µg/mL) when growing the AX2 VatM-GFP strain.

### Infection assays to detect SCVs by confocal microscopy

*D. discoideum* AX2 VatM-GFP grown in suspension axenically in HL5 medium was used. Amoebae were prepared by three cycles of centrifugation at 210 x *g* during 5 min at 4°C and resuspension in 1 mL of Soerensen buffer (2 g/L KH_2_PO_4_, 0.36 g/L Na_2_HPO_4_ x 2H_2_O, pH 6.0). Next, a suspension containing 1-2 x 10^6^ amoeba/mL was prepared after counting viable cells in a Neubauer chamber. Bacteria were prepared from overnight (O/N) cultures by centrifuging at 3,420 x *g* during 5 min at 4°C and suspended in Soerensen buffer. Amoebae were infected in a final volume of 0.2 mL using a multiplicity of infection (MOI) of 100 bacteria/cell and incubated at 23°C during 1 h without shaking to allow bacterial internalization. Next, infected amoebae were centrifuged at 210 x *g* during 5 min at 4°C, and the pellet was washed 3 times in Soerensen buffer to eliminate extracellular bacteria. Finally, infected amoebae were suspended in 0.2 mL of Soerensen buffer and incubated at 23°C for up to 4.5 h. Each sample was centrifuged at 210 x *g* during 5 min at 4°C and the pellet was suspended in 40 µL of Soerensen buffer. All samples were individually mounted on a glass slide on top of a thin layer (100 µL) of 1% agarose in PBS. A coverslip was placed over the sample and the borders sealed with colorless nail polish. Images were acquired with a Zeiss LSM 710 confocal microscope using the ZEN 2012 Back software (Zeiss). To detect the GFP label (amoebae), the sample was excited at 488 nm with an Argon laser and the emitted fluorescence was detected with a 493-549 nm filter. To detect mCherry label (bacteria), the sample was excited at 543 nm with a HeNe laser and fluorescence was detected with a 548-679 nm filter. All images were analyzed and edited using FIJI software (80, 81).

### Infection assays to evaluate intracellular survival

The infection procedure was performed as indicated above. Next, infected amoebae were suspended in 1 mL of Soerensen buffer and incubated at 23°C for up to 6 h. For each time point analyzed, an aliquot was obtained and used to determine viable amoebae by Trypan staining and counting in a Neubauer chamber. Another aliquot was used to determine titers of intracellular bacteria. To do this, infected amoebae were washed once with Soerensen buffer supplemented with 10 µg/mL gentamicin, centrifuged at 210 x *g* during 5 min, washed with Soerensen buffer to remove the antibiotic and centrifuged at 210 x *g* during 5 min.

Finally, the infected amoebae were lysed using 0.1% Triton X-100, serially diluted in PBS and plated on LB agar to determine CFUs. The internalization of each strain was calculated as intracellular CFUs after the hour of internalization (t=0) divided by the inoculated CFUs. The intracellular survival was calculated as the intracellular CFUs at 3 and 6 h post-infection (t=3 and t=6, respectively) divided by the intracellular CFUs at t=0. Statistical significance was determined by a two-way ANOVA with Dunnett’s test. All experiments were performed at least in biological triplicates.

### Isolation of highly enriched SCVs from infected amoebae

Infected *D. discoideum* AX2 cells were used to obtain a fraction enriched in vacuolar compartments containing *S*. Typhimurium according to the protocol described in (43), with modifications. Amoebae grown in T225 tissue culture flasks in HL5 medium were washed 3 times with Soerensen buffer. Bacteria from late exponential phase cultures were collected by centrifugation at 3,420 x *g* during 5 min at 4°C, and suspended in 50 mL of Soerensen buffer. Amoebae were infected using a MOI of 100 bacteria/cell and incubated at 23°C to allow bacterial internalization. In the case of the wild-type strain 1.2 x 10^8^ amoebae were infected, while in the case of the Δ*sopB* and Δ*sifA* mutants 2.4 x 10^8^ amoebae were infected. Next, extracellular bacteria were removed by washing 3 times with Soerensen buffer. Finally, infected amoebae were maintained in Soerensen buffer and incubated during 3 h at 23°C. After completion of the infection procedure, amoebae were washed 3 times with Homogenization Buffer (HB; 150 mM sucrose, 0.5 mM EGTA, 20 mM HEPES pH 7.4), scrapped from the tissue culture flask, and suspended in 4 mL of HB supplemented with cOmplete™ Protease Inhibitor Cocktail (Roche) and 5 µg/mL cytochalasin D (Sigma-Aldrich) (HB complete), to decrease organelle clumping. Each cell suspension was then transferred to a Dounce homogenizer (Sigma-Aldrich) and lysed by stroking the pestle 35-40 times (1 stroke = 1 up + 1 down) until more than 80% of free nuclei were visible under a light microscope. The homogenate was centrifuged at 100 x *g* during 5 min at 4°C to remove cell debris, the supernatant was collected, and the pellet suspended in 1 mL of HB complete and centrifuged again. This procedure was repeated 2 times and the supernatants from the 3 centrifugations were combined (∼6 mL) and defined as PNS. In the case of the PNS from non-infected control, 1 x 10^8^ CFU of the wild-type strain were added. Each PNS was loaded on top of a linear 10% (1.08 g/cm^3^) to 25% (1.15 g/cm^3^) OptiPrep gradient (Sigma-Aldrich) in HB with a 50% cushion prepared in 14 x 89 mm tubes (Beckman Coulter). Gradients were centrifuged at 210,000 x *g* during 3 h at 4°C using a SW-41 swinging bucket rotor (Beckman) in an Optima L-100 XP Ultracentrifuge (Beckman Coulter) with low acceleration and no brake settings. After centrifugation, 12 fractions of 1 mL were collected from the top of the gradient. Each fraction was analyzed by titrating the CFUs via serial dilution and plating, and by measuring the refraction index in a refractometer to determine its density.

### ELISA for *Salmonella* quantification

An anti-*Salmonella* ELISA developed to determine the presence of bacteria inside intact vacuoles was performed as described (43). A 96-well ELISA plate (Nunclon^®^ Immobilon) was coated with 70 µL of polyclonal rabbit anti-*Salmonella* antibody ab35156 (Abcam) suspended in PBS (5 µg/mL; 1:1,000) and incubated O/N at 4°C. The coated plate was blocked using 200 µL of blocking buffer (2% BSA from Sigma-Aldrich in PBS) during 1.5 h at room temperature. The plate was then washed four times with PBS and the different samples were loaded. To quantify bacteria per fraction, a standard curve of serial 2-fold dilutions of *S*. Typhimurium 14028s wild-type was prepared. For the fraction samples, an aliquot of 50 µL from F6 to F9 were diluted in 500 µL of HB (for pre-osmotic shock treatment) or in H_2_O (for osmotic shock treatment). Next, each sample was loaded on different wells of the ELISA plate and incubated during 1 h at room temperature. After that, the plate was washed with PBS and incubated O/N at 4°C with biotinylated rabbit anti-*Salmonella* antibody ab35156 (Abcam) diluted in blocking buffer (2 µg/mL; 1:2,000). Then, the plate was washed and incubated with Streptavidin-Peroxidase diluted in blocking buffer (Sigma-Aldrich, 1 mg/mL, 1:5000) during 1 h at room temperature. Subsequently, the plate was washed 6 times with PBS, the buffer was removed, 100 µL of SigmaFast OPD substrate solution (Sigma-Aldrich) was added, and the plate was incubated during 20-30 min at room temperature protected from light. The reaction was stopped by adding 50 µL of 10% SDS, and absorbance at 450 nm was determined in a FluoSTAR Omega microplate reader (BMG Labtech) at 450 nm.

### Protein precipitation and SDS-PAGE

Protein content in each fraction obtained was determined using the Micro BCA Protein Assay kit (Thermo Fisher Scientific) according to the manufacturer instructions. Then, each sample was precipitated using the chloroform/methanol method (82). To do this, a volume of sample containing 20 µg of protein was mixed thoroughly with 4 volumes of methanol, and then 1 volume of chloroform was added and mixed. After that, 3 volumes of milliQ water were added, the mixture was thoroughly vortexed and then centrifuged at 16,100 x *g* during 10 min at 4°C. The organic layer was discarded and 3 volumes of methanol were added, mixed thoroughly and centrifuged as mentioned. Finally, the supernatant was discarded and the protein pellet was air-dried and stored frozen at -20°C. The protein pellets corresponding to F6 and F7 from each experiment and condition were combined in 30 µL of Laemmli sample buffer (Bio-Rad) and then heated at 95°C during 10 min. The samples were loaded in a NuPAGE 4-12% Bis-Tris pre-cast Gel (Invitrogen) and run at 200 V constant during 45 min. The gel was then fixed during 30 min in an aqueous solution containing 10% acetic acid and 40% ethanol. Proteins were stained by O/N incubation in colloidal Coomassie blue G-250 (8% ammonium sulfate, 0.8% phosphoric acid, 0.08% Coomassie blue G-250, 20% ethanol) and destained in milliQ water until bands were visible.

### Mass spectrometry sample preparation

Each gel lane corresponding to a sample was cut into 10 pieces and destained in 100 µL of a 1:1 mixture of acetonitrile and 0.2 M ammonium bicarbonate pH 8. Each gel piece was incubated at 30°C during 30 min with shaking. The liquid was discarded and the process repeated one more time. Then, cysteines were reduced by incubating in 10 mM DTT in 67 mM ammonium bicarbonate pH 8 at 56°C during 1 h with shaking. The reduced cysteines were then alkylated using 55 mM iodoacetamide in 67 mM ammonium bicarbonate pH 8 and incubating at 25°C during 45 min with shaking. The gel pieces were then desiccated by adding 100% acetonitrile and incubating at 30°C during 30 min with shaking. In-gel protein digestion was performed by O/N incubation of the gel pieces with 1 µg of sequencing grade modified trypsin (Promega) in 67 mM ammonium bicarbonate pH 8 at 37°C. The next day, peptides from tryptic digestions were eluted from the gel pieces using a mixture containing 42.5% 50 mM ammonium bicarbonate, 42.5% acetonitrile and 5% formic acid, and incubating at 30°C during 1 h with shaking. Tryptic digestion samples were vacuum-dried and suspended in 2% acetonitrile, 0.1% formic acid in H_2_O (solvent A), and sonicated during 10 min in a water bath. The samples were then desalted using Bond Elute OMIX C18 tip filters (Agilent) according to the manufacturer instructions, and the peptides were finally eluted in 50% acetonitrile, 1% formic acid in H_2_O. These samples were vacuum-dried again and suspended in 10 µL of solvent A, sonicated during 10 min in a water bath. Finally, the peptide concentration of each sample was determined in a NanoDrop by measuring absorbance at 280 nm prior to its analysis by LC MS/MS.

### LC MS/MS data acquisition

Tryptic digests were analyzed by nano-LC MS/MS. To do this, each sample was injected into a nano-HPLC system (EASY-nLC 1000, Thermo Scientific) fitted with a reverse-phase column (Acclaim column, 15 cm x 50 µm ID, PepMap RSLC C18, 2 µm, 100 Å pore size, Thermo Scientific) equilibrated in solvent A. Peptides were separated at a flow rate of 300 nL/min using a linear gradient of 3% to 55% solvent B (80% acetonitrile, 0.08% formic acid) during 30 min. Peptide analysis was conducted in a Q-Exactive Plus mass spectrometer (Thermo-Scientific) set in data-dependent acquisition mode using a 30 s dynamic exclusion list. A resolution of 70,000 (at *m/z* 400) was used for MS scans. The 10 most intense ions were selected for HCD fragmentation and fragments were analyzed in the Orbitrap.

### Proteomic data analysis

A target-decoy database including sequences from *Dictyostelium discoideum* (taxon identifier: 44689), *Salmonella enterica* subspecies *enterica* serovar Typhimurium strain 14028s (taxon identifier: 588858) downloaded from Uniprot consortium in March 2018, and 127 most common mass spectrometry contaminants was generated using PatternLab for Proteomics version 4.0 (44). For protein identification, the Comet search engine was set as follows: tryptic peptides; oxidation of methionine and carbamidomethylation as variable modifications; and 40 ppm of tolerance from the measured precursor *m/z*. XCorr and Z-Score were used as the primary and secondary search engine scores, respectively. Peptide spectrum matches were filtered using the Search Engine Processor (SEPro) and acceptable false discovery rate (FDR) criteria was set on 1% at the protein level. The Approximately Area Proportional Venn Diagram module was used to perform comparisons between conditions and to determine proteins uniquely identified in each situation. Proteins found in at least three biological replicates of one condition were considered as “uniquely identified” when absent in all replicates of the other condition. For enrichment analysis, the TFold module was used to generate a volcano plot of the samples. This tool maximizes the identifications of proteins differentially detected between two conditions that satisfies both a Fold-change cutoff (that varies with the *t*-test P-value) and a stringency criterion that aims to detect proteins of low abundance under a Benjamini and Hochberg False Discovery Rate (FDR) estimator. For all comparisons, the FDR was fixed at 0.05. The TFold module then explored several values of the F-Stringency parameter and selected the one that maximizes the number of differentially detected proteins between the two datasets, for the specified Q-value of 0.05 (83, 84). Functional clustering analyses were performed using DAVID (45, 46). The mass spectrometry proteomics data have been deposited to the ProteomeXchange Consortium via the PRIDE (85) partner repository with the dataset identifier PXD014955.

## Acknowledgments

This work was supported by FONDECYT grants 1140754 and 1171844, to CAS. CV and ÍU were supported by CONICYT fellowships 21140615 and 21150005, respectively. The team of JE is supported by an ERC CoG grant (EndoSubvert), and acknowledges support from the ANR (StopBugEntry and AutoHostPath programs). JE is member of the LabExes IBEID and Milieu Interieur. The funders had no role in study design, data collection and interpretation, or the decision to submit the work for publication. We thank Dr. Macarena Varas and Carolina Hernández (Unidad de Microscopía Confocal, Facultad de Ciencias Químicas y Farmacéuticas, Universidad de Chile) for their technical assistance in microscopy experiments. We are grateful to Dr. Magalie Duchateau and Dr. Mariette Matondo (Mass Spectrometry for Biology - UTechS MSBio, Institut Pasteur) for their technical assistance in nano-LC MS/MS analyses.

## Author Contributions

Conceptualization: CV, MG, JE and CAS; methodology: CV, MG, IMU, AS, JE and CAS; resources: CV, MG, JE and CAS; investigation: CV, MG, IMU and AS; formal analysis: CV and MG; supervision: JE and CAS; project administration: CV, MG, AS and CAS; funding acquisition: CV and CAS; visualization: CV and MG; writing-original draft preparation: CV, MG, JE and CAS; writing-review and editing: CV, MG, IMU, AS, JE and CAS. All authors read and approved the final manuscript

